# Melanocortin receptor 4 agonist setmelanotide treats opioid-induced respiratory depression

**DOI:** 10.64898/2026.03.08.708886

**Authors:** Mateus R. Amorim, Noah R. Williams, Melanie A. Ruiz, Junia L. de Deus, O Aung, Olga Dergacheva, Joan Escobar, Mi-Kyung Shin, Cole R. Winston, Thales H. C. Furquim, Jeffrey S. Berger, David Mendelowitz, Vsevolod Y. Polotsky

## Abstract

**Background:** The primary cause of death associated with opioids is opioid-induced respiratory depression (OIRD). Naloxone is used to reverse OIRD, but this drug is a competitive antagonist of µ-opioid receptor (MOR) and reverses analgesia, which limits its therapeutic use. Alternative non-opioid receptor antagonist-based approaches to OIRD treatment and prevention are needed. The aim of this study was to evaluate if setmelanotide (SET) is capable of reversing OIRD in a mouse model.

**Methods:** C57BL/6J male and female mice and Sprague-Dawley rats were given IP morphine or fentanyl and then treated 15 min later with either SET or vehicle VEH (IP) in a random order. Breathing was recorded by barometric plethysmography, and pain sensitivity was measured by the tail-flick test.

**Results:** In mice with OIRD, SET induced a 3-fold reduction of the apnea index, and decreased apnea duration as compared to the VEH treatment. SET increased respiratory rate and did not affect opioid-induced analgesia. Photostimulation of MC4R+ ChR2-expressing fibers in the parafacial region of MC4R-Cre mice elicited short-latency excitatory postsynaptic current in rostral ventral respiratory group (rVRG) pre-motoneurons projecting to the phrenic nucleus in the C3-C4 ventral horns of the spinal cord. Fentanyl inhibited the activity of rVRG neurons and SET reversed this effect.

**Conclusions:** SET effectively treated OIRD by increasing respiratory rate and inducing a significant decrease in the number of apneas without decreasing analgesia.

## Introduction

Drug overdose has become a national crisis in the US with drug overdose deaths increasing from 16,849 in 1999 to 106,699 in 2021^1^. The majority of these deaths are related to opioids, rising from 8,048 in 1999 to 80,411 in 2021^1^. The primary cause of death associated with opioids is opioid-induced respiratory depression (OIRD), characterized by reduced respiratory rate, decreased airflow, and central apneas, resulting in severe hypoxemia and cardio-respiratory arrest^2,3^. There are well-defined patient categories at high risk of OIRD that would benefit from acute OIRD prevention. Obesity predisposes to chronic pain and increases the risk of opioid use disorder^4^, increasing significantly the risk of opioid-induced mortality^5–7^.

Opioids act on μ, Δ and κ opioid receptors expressed in the brain^8^. Opioid binding to μ-opioid receptor (MOR), a G protein-coupled receptor, alleviates pain through G protein signaling, and induces OIRD *via* β-arrestin activation^3,9^. Naloxone rapidly reverses OIRD, however, it has short half-life (∼ 1 hr) blocks the analgesic effects of opioids^10^ and causing acute withdrawal, limiting its use^11,12^. Thus, MOR antagonist alternatives are needed for OIRD treatment.

Non-opioid respiratory stimulants have been considered as a potentially viable alternative to MOR antagonists for OIRD treatment and prevention. The candidates include potassium channel blockers, N-methyl-D-aspartate receptors antagonists, ampakines and analeptics. However, proof-of-concept clinical trials showed either low efficacy or significant adverse effects^13,14^.

We have shown that leptin, an adipocyte-produced hormone, stimulates breathing by increasing the hypercapnic ventilatory response (HCVR) and treats hypoventilation in obese animals^15–17^. We have also shown that leptin prevents OIRD and decreases opioid mortality in mice^18,19^. However, leptin resistance often limits leptin as a pharmacotherapy^20–23^. Leptin upregulates pro-opiomelanocortin (POMC)^24–27^, which is a pre-hormone post-transcriptionally processed into several peptides, including α-melanocyte stimulating hormone (α-MSH), a ligand for the melanocortin 4 receptor (MC4R)^28–31^. The MC4R agonist setmelanotide (SET) has been approved by the FDA for treatment of genetic forms of obesity^32–35^. We have previously shown that SET increases minute ventilation and effectively treats sleep disordered breathing (SDB) in diet-induced obese (DIO) mice^36^. Furthermore, we have shown that respiratory effects of MC4R agonists act *via* CO_2_/H+ sensitive neurons of the retrotrapezoid nucleus (RTN) and that MC4R (+) RTN neurons project to brainstem respiratory premotor neurons, which are connected to phrenic motoneurons in the cervical spinal cord.

In the present study we hypothesized that SET may treat OIRD, without affecting analgesia. We challenged lean and diet-induced obese C57BL/6J mice with high doses of morphine and fentanyl and then examined if SET administered after morphine and fentanyl reversed OIRD. We also examined the effect of SET on analgesia by the tail flick test and we also explored potential mechanisms of MC4R agonists in OIRD *ex vivo* by optogenetic approach in *Mc4r-Cre* mice probing if photoexcitation of MC4R+ RTN neurons induces excitatory neurotransmission to brainstem respiratory premotor neurons and by determining SET affinity for MOR in HEK-293 cells. Finally, we examined the effects of SET and fentanyl on excitatory postsynaptic currents (EPSCs) in the brainstem respiratory premotor neurons.

## Material and Methods

Experiments were conducted in a total of 101 animals, including lean and diet-induced obese (DIO) C57BL/6J male, DIO female mice, lean Sprague Dawley rats, and *Mc4r*-*Cre* mice. Animals were housed under controlled temperature (29°C) and a 12-hour light/dark cycle, with free access to water and food. Depending on the experimental group, they were fed either a standard chow or a high-fat diet. All experimental procedures were approved by The George Washington University (Protocol # A2023-014).

To assess breathing, animals underwent whole-body plethysmography (WBP) to measure respiratory rate, minute ventilation, and apnea index. A two-phase crossover study was used, involving intraperitoneal administration of opioids (morphine 10 mg/kg or fentanyl 0.5 mg/kg, IP) in combination with setmelanotide (SET; 1 mg/kg in 2% heat-inactivated mouse serum plus 0.5% DMSO) or vehicle (VEH; mouse serum plus 0.5% DMSO). Measurements were performed in calibrated chambers using high-precision signal acquisition systems. Animals were acclimated to the apparatus before testing, and breathing was recorded after drug administration over a two-hour period.

Analgesic effects were assessed using the tail-flick test in response to a thermal stimulus with hot water, measuring tail withdrawal at 15, 30, 60, and 120 minutes after administration of SET or VEH.

Electrophysiological analyses were done on rVRG MC4R+ neurons that project to the phrenic nucleus in the spinal cord. Whole-cell patch-clamp recordings were obtained in voltage-clamp configuration to measure postsynaptic excitatory currents (EPSCs) *in vitro*. Changes in the frequency and amplitude of EPSCs were analyzed after fentanyl and SET administration. SET binding to MOR was assessed in the agonist radioligand Receptor Binding Assay (Eurofins Cerep, Celle l’Evescault, France).

Data analyses were done using Prism software, version 10.4.1 (GraphPad Software Inc., San Diego, CA, USA). To assess the effects of experimental factors and potential interactions, we applied a mixed-effects model. Depending on the data structure, comparisons within groups were conducted using two or one-way ANOVA, paired t-tests, or Wilcoxon matched-pairs signed-rank tests. Differences between groups were evaluated with the Mann–Whitney test. Statistical significance was defined as *P* ≤ 0.05. Descriptive statistics were extracted from the summary outputs of the corresponding analyses and are reported as means ± standard error. For graphical representation, data were displayed as boxplots with individual animal values showing individual mice.

Additional methodological details are provided in the online data supplement.

## Results

### 1. SET increased respiratory rate and reduced apnea index in lean and DIO male mice challenged with morphine without affecting analgesia

To examine whether SET improves breathing in our OIRD mouse model, we performed the breathing recordings in a crossover, randomized manner (Figure 1 A). In lean male mice, morphine significantly increased the number of apneic events and respiratory rate (RR) (Figure 1C). A single dose of SET administered 15 minutes after morphine significantly stimulated breathing, as evidenced by an increase in RR (P < 0.05) and reduced apnea index (P < 0.05) (Figure 1 B – C). The longitudinal breath-by-breath analysis showed that morphine induced a significant increase in the apnea frequency throughout the entire 2-hour recording and SET attenuated opioid-induced apneas. In DIO male mice, morphine reduced the RR and increased apnea index, as observed in our previous study^18^. Importantly, SET increased RR and decreased apnea frequency and duration (P < 0.05) (Figure 1 D and E). Compared to baseline, morphine led to breathing instability, as indicated by higher values of short-term (SD1) and long-term (SD2) V_E_ variability in the Poincaré analysis in both lean and DIO male mice. Obesity was associated with higher breathing instability after morphine administration, as DIO mice exhibited greater SD1 (0.23 ± 0.06 vs 0.15 ± 0.03, *P* = 0.0011) and SD2 variability (1.57 ± 0.48 vs 0.73 ± 0.15, *P* = 0,0011) in comparison to lean mice. SET treatment did not stabilize breathing in either lean or DIO mice (Figure 2). Thus, a single dose of the MC4R agonist SET stimulated control of breathing and improved upper airway patency in OIRD.

**Figure 1:**
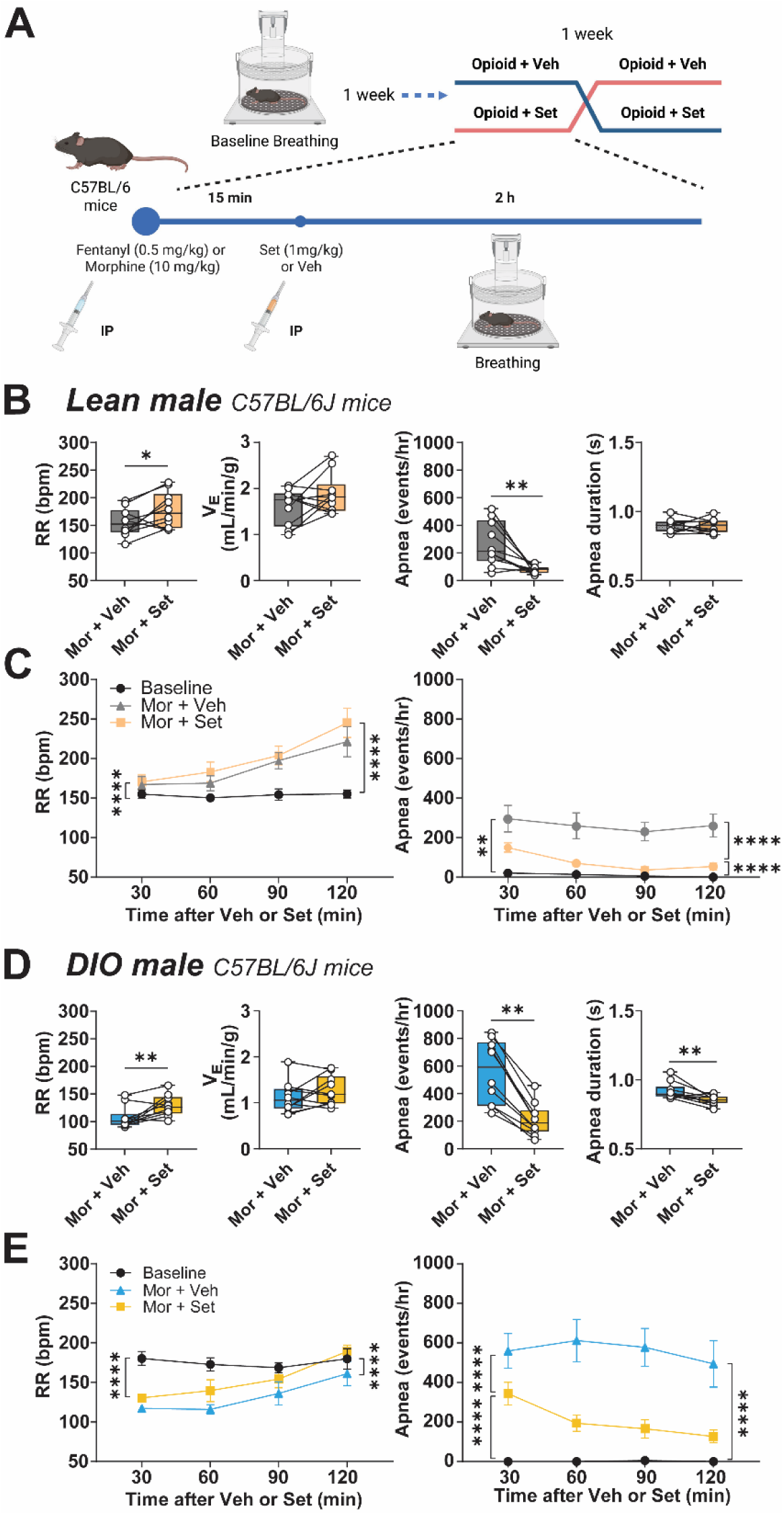
Setmelanotide (Set) administered 15 minutes after morphine reduced the apnea index and increased respiratory rate (RR) in lean and diet-induced obese (DIO) mice. **(A)** Experimental design. (B) Individual and grouped data showing the differences between the morphine + vehicle (Mor + Veh) and Mor + Set effects on the respiratory rate (RR), minute ventilation (V_E_), apnea index and duration in lean male (N = 10) and (D) DIO male (N = 10). * *P* ≤ 0.05 and, ** *P* < 0.01 Wilcoxon matched-pairs signed rank test, upper panels. (C) Time course of RR and apnea index throughout 2-hour recording at baseline (N = 7), Mor + Veh, and Mor + Set in lean, and (E) DIO male mice (baseline, N = 7). Means ± SEM. ** *P* < 0.01 and **** *P* < 0.0001 effect of treatment using two-way ANOVA.

**Figure 2:**
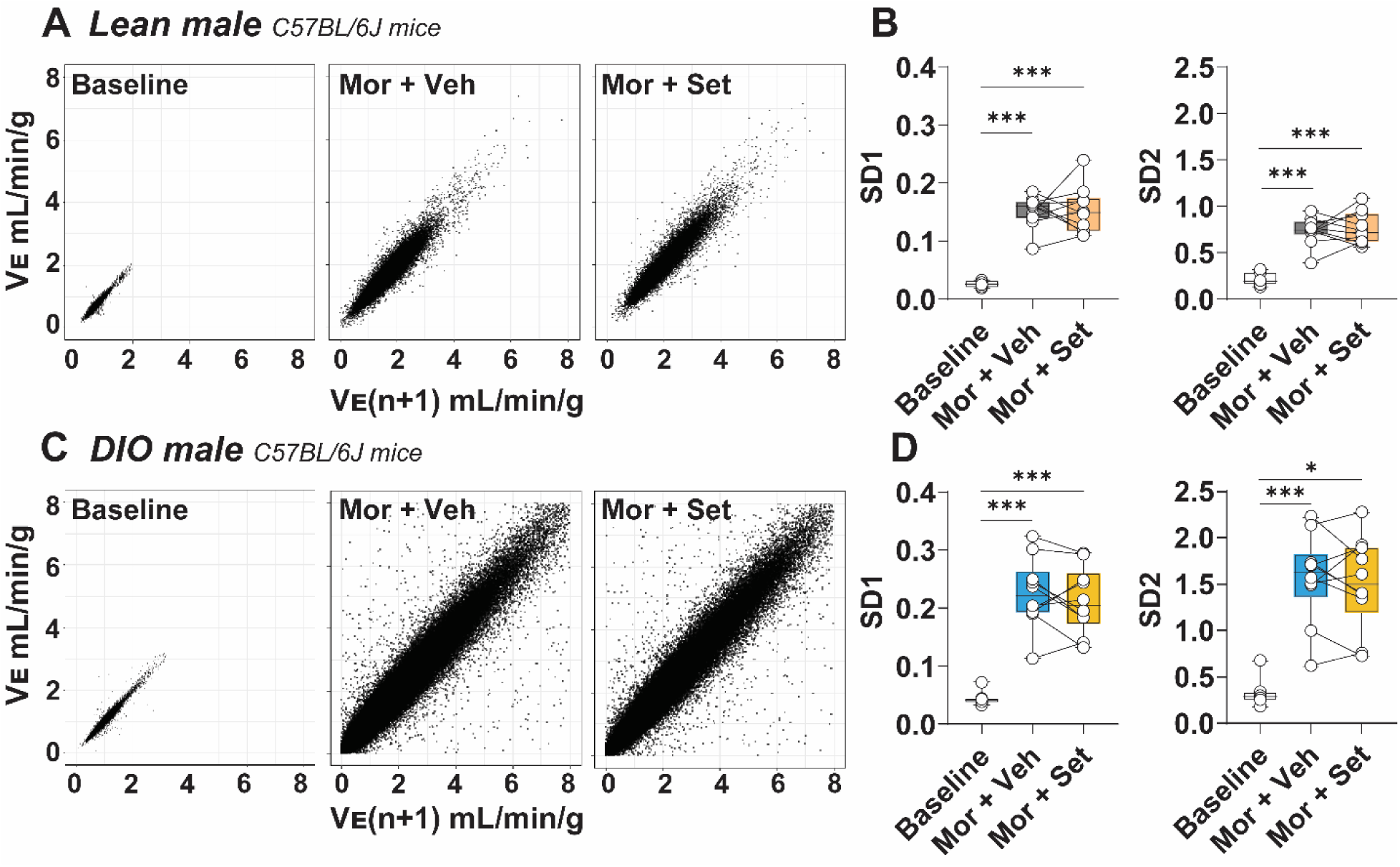
Setmelanotide (Set) did not attenuate morphine (Mor)-induced ventilatory instability in lean and diet-induced obese (DIO) male mice. **(A)** Representative Poincaré plots of minute ventilation (V_E_) in lean C57BL/6J male mice at baseline, after morphine plus vehicle (Mor + Veh), and after Mor plus setmelanotide (Mor + Set) administration. **(B)** Short-term (SD1) and long-term (SD2) breathing variability in lean male mice. **(C)** Representative Poincaré plots of V_E_ from lean C57BL/6J diet-induced DIO male mice at baseline, Mor + Veh, and after Mor + Set administration. **(D)** SD1 and SD2 breathing variability in DIO male mice. Data are presented as median ± IQR. Statistical analyses were performed using Mann–Whitney test. * *P* ≤ 0.05 and *** *P* < 0.001.

The potential of SET to enhance breathing without impairing analgesia was examined using the tail-flick test, conducted in a randomized crossover design (Figure 3A). As expected, we observed an increase in tail-flick latency after morphine administration in both lean and DIO male mice (Figure 3 B and C). SET did not significantly affect tail-flick latency measured every 15 minutes after morphine bolus, and this observation was consistent in both lean and DIO male mice (Figure 3 B and C).

**Figure 3:**
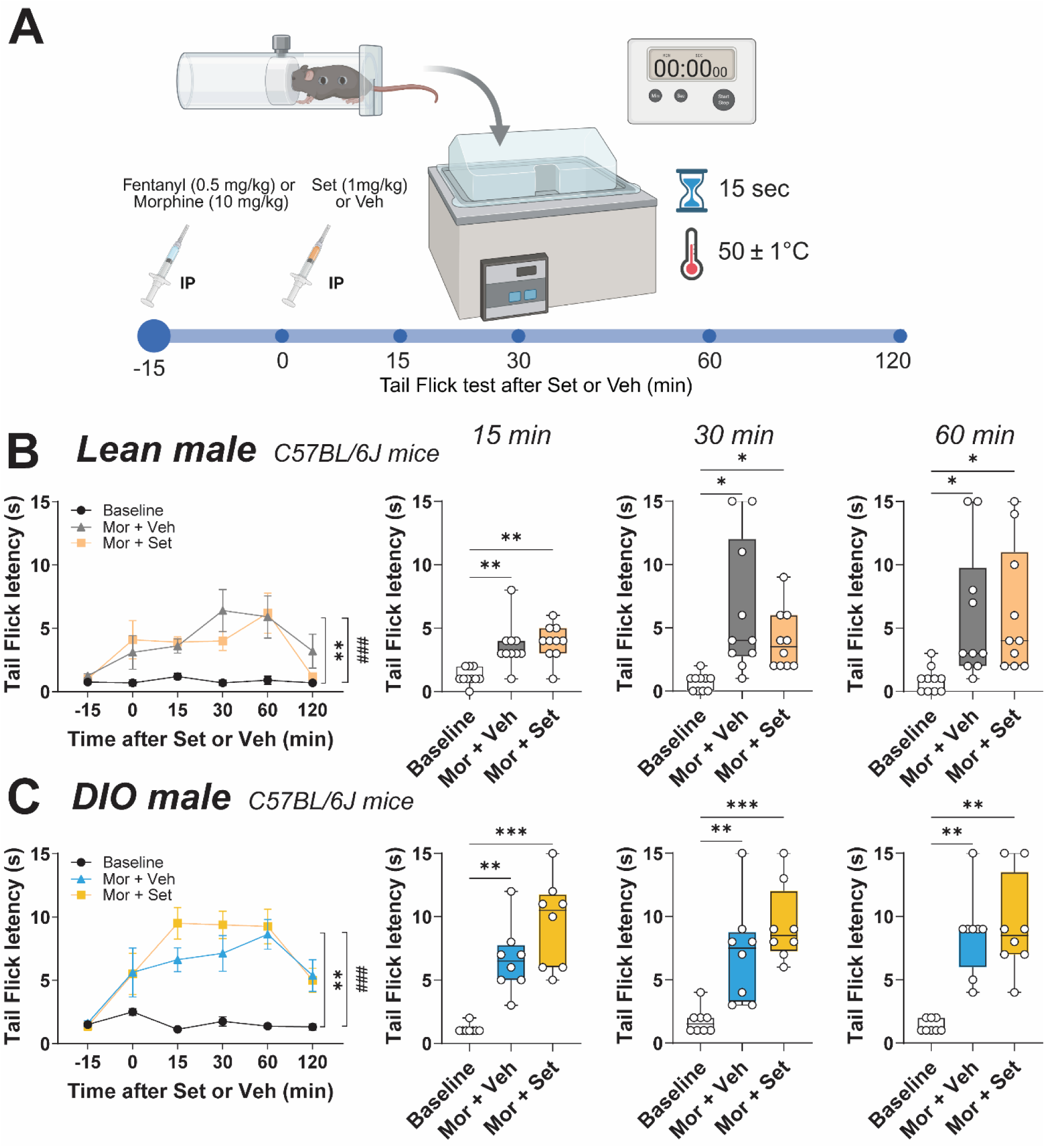
Setmelanotide (Set) administered 15 minutes after morphine did not affect morphine-induced analgesia. **(A)** Experimental design. Tail-flick latencies were recorded after immersion of the distal third of the tail in a 50±1°C water bath. A maximum of 15 seconds immersion time was used to avoid tissue damage. Intraperitoneal (IP) injection of morphine (Mor) was followed by IP Set or vehicle (Veh) 15 minutes later. Tail-flick latencies were measured at baseline, before Set/vehicle treatment (0), and at 15, 30, 60, and 120 minutes after Set/vehicle. **(B,** left) Time course of tail-flick latency for baseline, Mor + Veh, and Mor + Set showing means ± SEM in lean male mice (N = 10), and **(C,** left) DIO male mice (N = 8). ** *P* < 0.01 and ^###^ *P* < 0.001 effect of treatment using two-way ANOVA. Morphine increased tail flick latency at 15, 30 and 60 minutes after morphine administration in **(B,** right) lean male mice, and **(C,** right) DIO male mice. Values are shown for individual mice. * *P* ≤ 0.05, ** *P* < 0.01 and **** *P* < 0.001 effect of treatment using repeated measures ANOVA followed by Tukey’s multiple comparisons test.

### 2. SET increased respiratory rate and reduced apnea index in lean and DIO male mice challenged with fentanyl without affecting analgesia

Next, we tested if SET attenuates OIRD in mice treated with fentanyl. We compared SET to VEH in lean and DIO male mice, examining RR, V_E_, apnea index, and duration. Fentanyl caused a marked reduction in RR and a significant increase in apnea index in both lean and DIO male mice. SET blunted the OIRD in both groups, significantly increasing RR and reducing the apnea index throughout the entire 2-hour recording period (P < 0.05) (Figure 4 A – D). To investigate whether these effects were sex-dependent, we also evaluated breathing in DIO female mice that received fentanyl followed by SET. In contrast to DIO males, RR was not significantly affected by fentanyl in DIO females (compare 4 D and F), the apnea index was increased. SET decreased the incidence of apneas in female mice (P < 0.05) (Figure 4 E and F). The Poincare analysis showed that fentanyl induced breathing instability, as reflected by increased SD1 and SD2 ventilatory variability in all groups of mice (Figure 5). In lean male mice, a single dose of SET blunted fentanyl-induced short-term ventilatory variability. Obesity leads to lower fentanyl-induced breathing instability, evidenced by decreased SD1 (0.08 ± 0.02 vs 0.15 ± 0.04, *P* = < 0.0001) and SD2 (0.36 ± 0.11 vs 0.71 ± 0.23, *P* = 0,0008) in relation to lean. In contrast, SET did not attenuate fentanyl-induced ventilatory instability in DIO male and female mice of both sexes after fentanyl administration (Figure 5 E and F). These findings suggest that the respiratory stimulant effects of SET in our mouse model of OIRD are both weight and sex-dependent.

**Figure 4:**
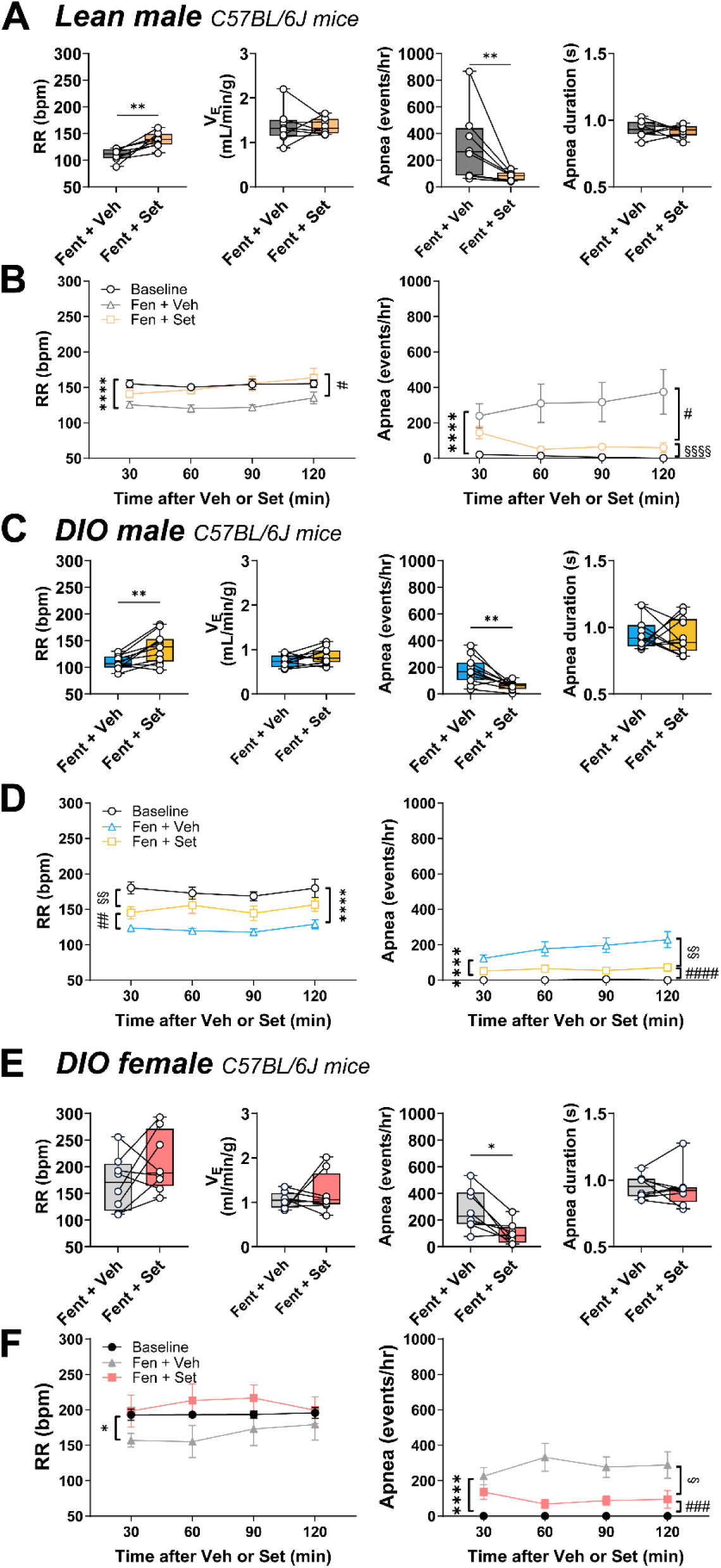
Setmelanotide (Set) reduced the apnea index and increased respiratory rate (RR) in lean and diet-induced obese (DIO) mice treated with fentanyl. **(A)** Individual and grouped data showing the differences between the fentanyl + vehicle (Fent + Veh) and Fent + Set effects on the respiratory rate (RR), minute ventilation (V_E_), apnea index and duration in lean male (N = 8), **(C)** DIO male (N = 11), and **(E)** DIO female (N = 8). * *P* ≤ 0.05 and, ** *P* < 0.01 Wilcoxon matched-pairs signed rank test. **(B)** Time course of RR and apnea index throughout 2-hour recording at baseline (N = 7), Fent + Veh, and Fent + Set in lean male, **(D)** DIO male mice (baseline, N = 7), and **(F)** DIO female mice (baseline, N = 5). Means ± SEM. * *P* ≤ 0.05, ** *P* < 0.01, *** *P* < 0.001 and **** *P* < 0.0001 effect of treatment using two-way ANOVA. The baseline group is the same as in Figure 1 for comparison purposes.

**Figure 5:**
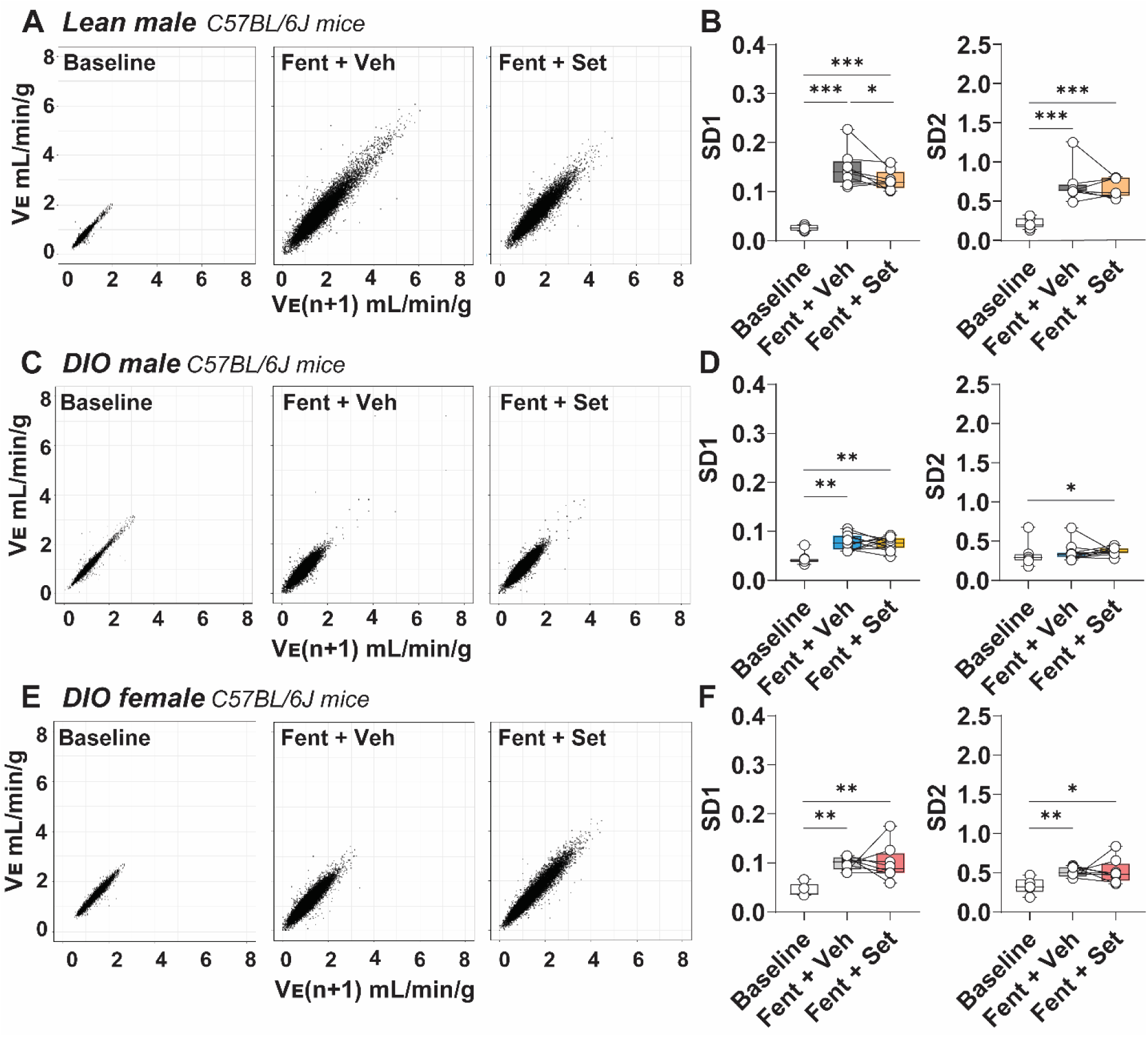
Setmelanotide (Set) attenuated fentanyl (Fent)-induced ventilatory instability in lean male mice. **(A)** Representative Poincaré plots of minute ventilation (V_E_) from lean C57BL/6J male mice at baseline, after Fent plus vehicle (Fent + Veh), and after Fent plus setmelanotide (Mor + Set) administration. **(B)** Short-term (SD1) and long-term (SD2) breathing variability in lean male mice. **(C)** Representative Poincaré plots of V_E_ from lean C57BL/6J diet-induced obese (DIO) male mice at baseline, Fent + Veh, and after Fent + Set administration. **(D)** SD1 and SD2 breathing variability in DIO male mice. **(E)** Representative Poincaré plots of V_E_ from lean C57BL/6J DIO female mice at baseline, Fent + Veh, and after Fent + Set administration. **(F)** SD1 and SD2 breathing variability in DIO female mice. Data are presented as median ± IQR. Statistical analyses were performed using Wilcoxon matched-pairs signed rank and Mann–Whitney tests. * *P* ≤ 0.05, ** *P* < 0.01 and *** *P* < 0.001. The baseline group is the same as in Figure 2 for comparison purposes.

As with morphine, fentanyl administration increased tail-flick latency in lean and DIO male mice, as well as in DIO female mice (Figure 6). SET did not attenuate fentanyl-induced antinociception at any of the time points evaluated.

**Figure 6:**
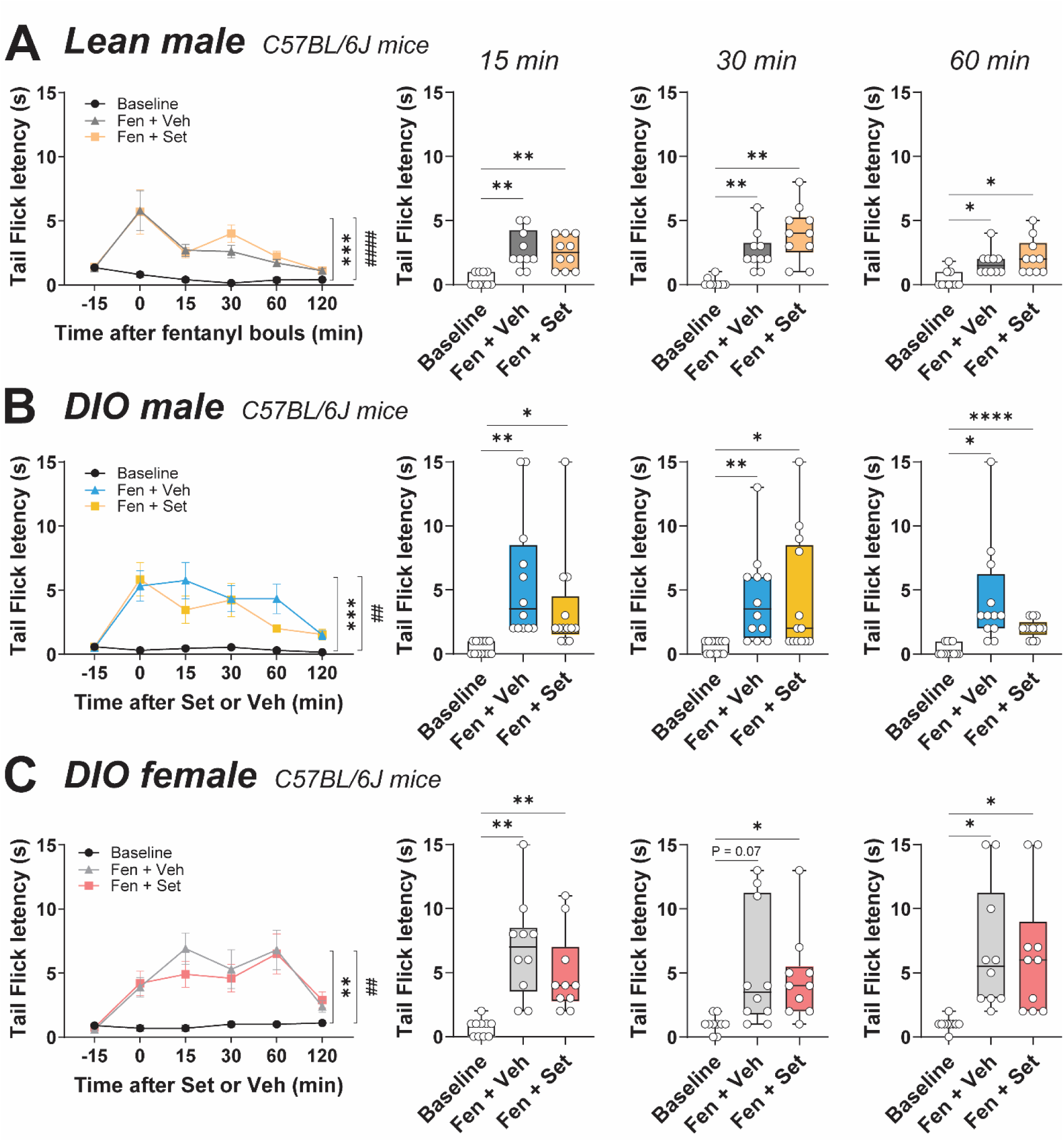
Setmelanotide (Set) did not affect fentanyl (Fent)-induced analgesia. **(A, left)** Time course of tail-flick latency for baseline, Fent + Veh, and Fent + Set showing means ± SEM in lean male mice (N = 10), **(B,** left) DIO male mice (N = 13) and **(C,** left) DIO female mice (N = 10). ** *P* < 0.01, *** *P* < 0.001 and ^###^ *P* < 0.001 effect of treatment using two-way ANOVA. Fentanyl increased tail flick latency at 15, 30 and 60 minutes after fentanyl administration in **(A,** right) lean male mice, **(B,** right) DIO male mice, and **(C,** right) DIO female mice. Values are shown for individual mice. * *P* ≤ 0.05, ** *P* < 0.01 and **** *P* < 0.001 effect of treatment using repeated measures ANOVA followed by Tukey’s multiple comparisons test.

### 3. SET induced reduction of the apnea index and decreased apnea duration in rats treated with fentanyl

In order to further explore translational relevance of our findings we examined the effect of SET on OIRD in another rodent model, rats. Following fentanyl administration (0.5 mg/kg IP) SET reduced the incidence of apneas, increased respiratory rate (134 ± 13 *vs* 94 ± 5 breaths/min in the vehicle group, P < 0.01), and more than doubled V_E_ (297 ± 44 *vs* 134 ± 12 ml/min, P < 0.01) (Figure 7). Altogether, these data are consistent with the notion that SET treated fentanyl-induced OIRD in rats, similar to benefits that occurred in mice.

**Figure 7:**
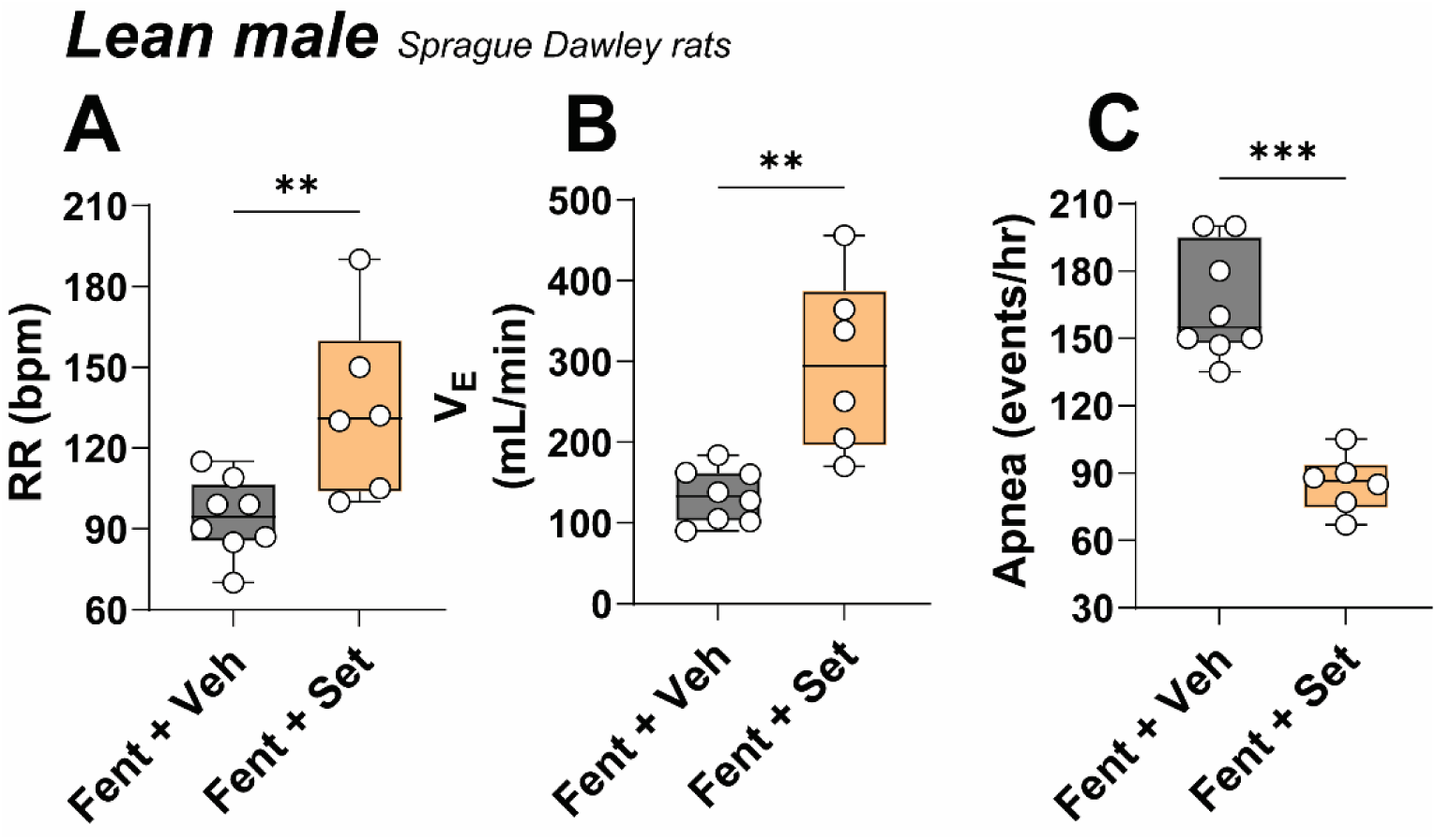
Setmelanotide (Set) increased respiratory rate (RR) and reduced apnea index in rats. V_E_, minute ventilation. Veh, vehicle (N = 8) and Set (N = 6). ** *P* < 0.01 and *** *P* < 0.001 effect of treatment using Mann Whitney test.

### 4. Parafacial MC4R+ neurons projects to respiratory pre-motor neurons, fentanyl inhibits rVRG neurons and SET reverses fentanyl inhibition

RTN neurons do not express MORs and are not sensitive to opioids^37,38^. RTN neurons do not project directly to the spinal cord, but have projections to the critical and opioid vulnerable downstream rVRG neurons^39,40^, which express MORs, and are inhibited by opioids^41^. rVRG neurons project to and excite neurons in the phrenic nucleus of the spinal cord. To express ChR2 in *Mc4r-Cre* RTN neurons mice were injected with *Cre*-dependent ChR2 virus into the RTN and to label rVRG neurons that project to the phrenic motor nucleus the C3/4 segments of the spinal cord were injected with the retrograde tracer Cholera Toxin B (CTB) (Figure 8 A and B). Extensive projections of MC4R+ RTN neurons to retrogradely labelled CTB+ rVRG neurons were detected (Figure 8 C and D). Spontaneous and ChR2 evoked EPSCs were recorded in pre-motor rVRG neurons that project to the phrenic nucleus of the spinal cord. Photoexcitation of ChR2+ MC4R+ fibers surrounding rVRG neurons evoked EPSCs with short (∼5 msec) latencies in rVRG neurons (Figure 8 E and F). Photoactivation of ChR2+ fibers from MC4R+ RTN neurons excite rVRG neurons that, in turn, project to and provide excitation to the phrenic motor nucleus in the C3/4 region of the spinal cord.

**Figure 8:**
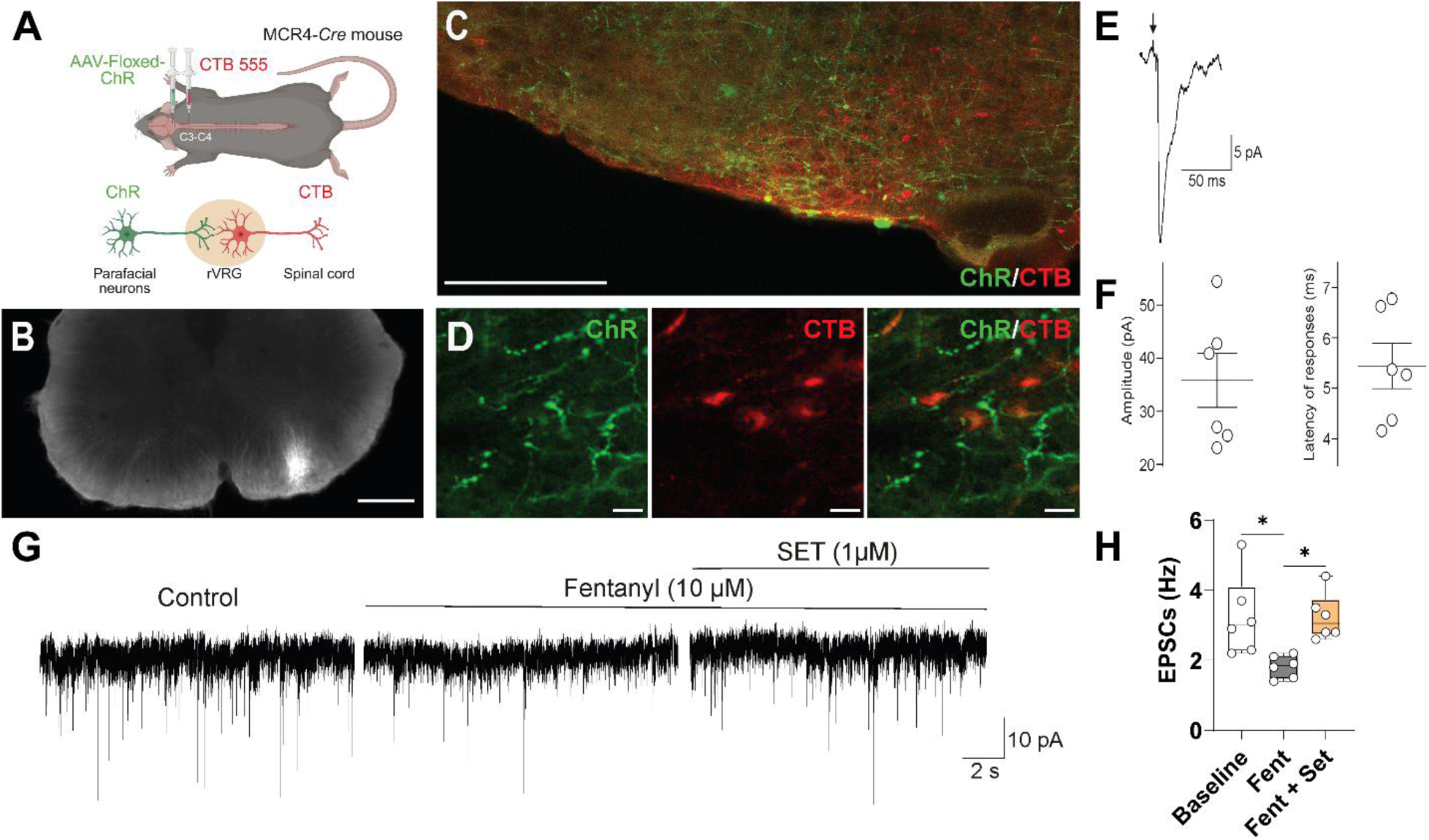
Parafacial MC4R+ neurons projects to respiratory pre-motor neurons. Fentanyl inhibits rostral ventral respiratory group (rVRG) neurons and setmelanotide (Set) reverses fentanyl inhibition. **(A)** *Cre*-dependent ChR2-EYFP was deployed in the parafacial region of *Mc4r-Cre* mice and the C3-C4 was injected with cholera toxin B (CTB)-Alexa 555. **(B)** Coronal slice of the spinal cord at the C3-C4 level showing the CTB injection site (white) in the ventrolateral area where phrenic motorneurons are densely located. Injection tract leaving the dorsal surface is also shown. Scale bar = 500 µM **(C)** Lower power image shows extensive density of MC4R+ green fibers surrounding CTB+ red cells. **(D)** Higher power images of the same region show (from left to right) MC4R+ fibers, CTB+ neurons and a merged image showing that green MC4R+ fibers enmesh the red CTB+ neurons (lower panels). Scale bar = 100 μm (upper panel) and 50 μm (lower panels). **(E)** Excitatory postsynaptic currents (EPSC) in a pre-motor rVRG neuron evoked by photostimulation of ChR2-EYFP-expressing MC4R fibers. The EPSCs were blocked at the end of experiment by the glutamate receptor antagonists AP-5 (50 μM) and CNQX (25 μM). **(F)** Summary data from 6 pre-phrenic motor neurons. Circles represent average from 80 sweeps in individual pre-motor neurons (3 ms stimulation at 1 Hz) while horizontal bars are population means ± SEM. Amplitude and latency of the glutamatergic responses are shown in the center and right, respectively. **(G)** Representative recordings of EPSCs in rVRG neurons that project to the phrenic nucleus at control conditions, after fentanyl followed by SET application to the rVRG neurons; **(H)** EPSCs frequency. n = 6 per group. * *P* ≤ 0.05.

Next, we tested the hypothesis whether MC4R agonists treat OIRD by stimulating downsteam respiratory premotor rVRG neurons that are inhibited by fentanyl. Our data shows that application of fentanyl caused a significant decrease in the frequency of glutamatergic EPSCs in rVRG pre-motoneurons directly projecting to the phrenic motor nucleus. Remarkably, photoexcitation of MC4R+ neurons reversed the inhibition of EPSCs in rVRG neurons induced by fentanyl (Figure 8 G and H).

Finally, we examined if SET may act directly binding to MOR. The competing radioligand binding assay showed that SET IC_50_ > 1.0^E-06^ M, the highest concentration tested, which indicated that SET has no affinity for MOR (Figure 9).

**Figure 9:**
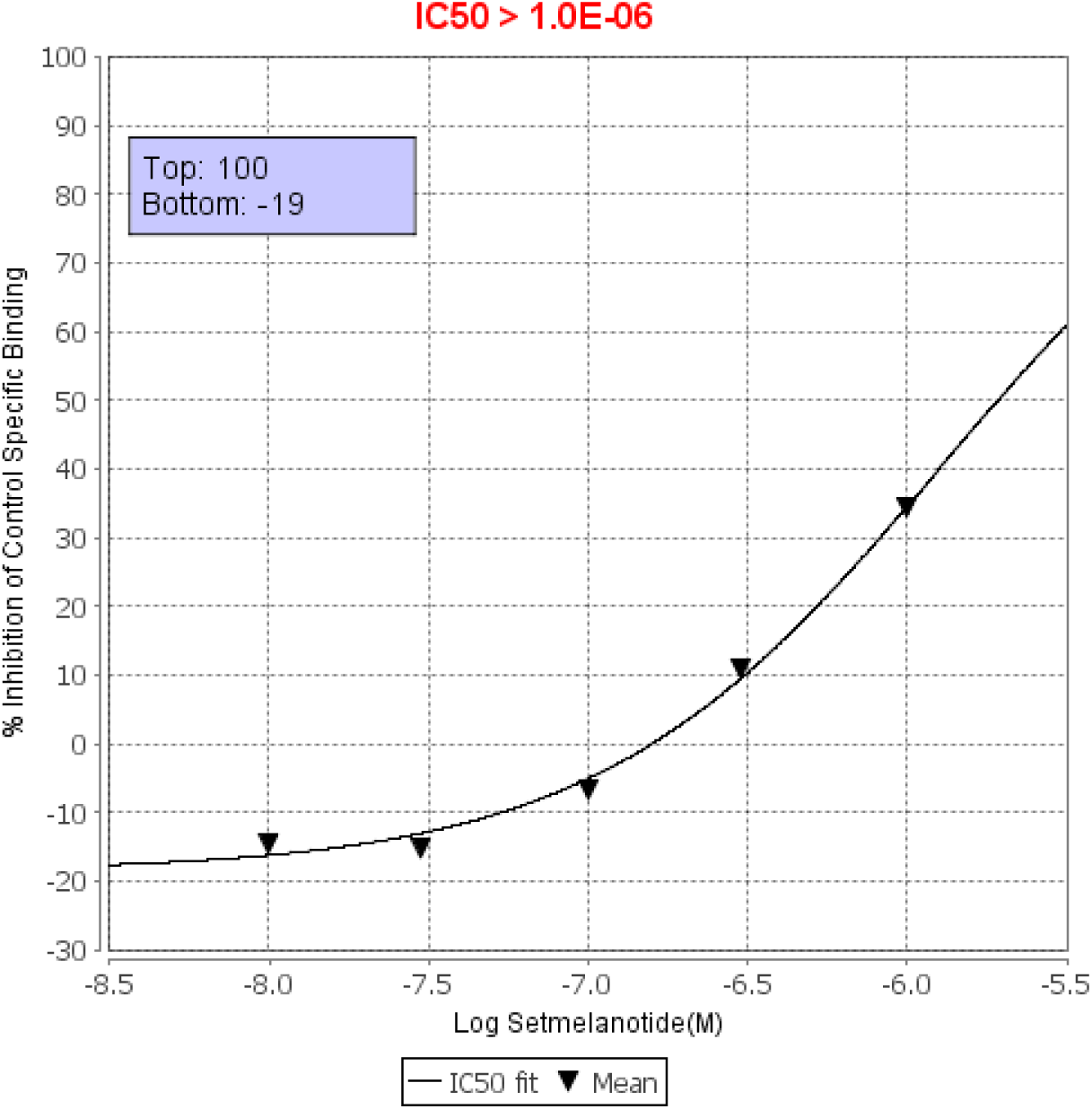
Setmelanotide on μ (MOP) (h) (agonist radioligand)

## Discussion

Our study provided the first evidence that a MC4R agonist SET effectively treats OIRD in rodent models. We have shown that SET reversed OIRD in both DIO mice with underlying SDB, in lean mice and rats, and that SET is effective in both morphine and fentanyl-induced OIRD without any effect on analgesia. We have identified rostral ventral respiratory group neurons (rVRG), which innervate phrenic motor neurons, as a potential target for SET. Finally, we have shown that SET has no affinity for MOR, but photoexcitation of MC4R+ neurons in the RTN induces excitatory postsynaptic currents (EPSCs) in rVRG neurons, which suggests a novel mechanism by which SET may reverse OIRD.

We have previously shown that A^y^ mice overexpressing the natural MC4R blocker AgRP hypoventilate during sleep and have suppressed hypercapnic ventilatory response (HCVR) compared to the weight-matched wildtype control mice^42^. Bassi *et al* showed that intracerebroventricular injections of the MC4R blocker SHU9119 decreased ventilatory responses to 8% CO_2_ and attenuated leptin-induced hyperventilation in rats^43^. Rivera et al. has shown that SET increases the hypercapnic ventilatory response^44^. We have shown that SET significantly increased HCVR and alleviated sleep disordered breathing (SDB) in DIO mice by increasing minute ventilation and decreasing a number of apneas during sleep^36^. Our present data in conscious, unanesthetized mice and rats show that SET effectively treated OIRD caused by morphine and fentanyl increasing respiratory rate and decreasing incidence of apneas without any effect on opioid analgesia, unlike MOR antagonists, naloxone and nalmefene. Obesity causes SDB and predispose to poor outcomes in OIRD^45^. We have shown that SET reverses OIRD in both obese and lean states. SET efficacy in two different species, mice and rats, emphasizes the robustness of our data.

Respiratory stimulants commonly act by enhancing the chemoreflex. Our recent data showed that *Mc4r* mRNA is colocalized with the paired-like homebox 2B transcription factor (*Phox2b*)^36^, which is universally present in CO_2_ sensing neurons, in two brainstem regions of ventilatory control, the nucleus of the solitary tract (NTS) and the retrotrapezoid nucleus (RTN)^37,39,46–48^, and neuromedin B (*Nmb*) ^36^, a highly specific marker of RTN CO_2_/H+ sensing neurons^49,50^. Our previous experiments in *Mc4r-Cre* mice transfected with *Cre*-dependent designer receptor exclusively activated with designer drugs (DREADD) showed that chemogenetic stimulation of MC4R+ neurons in RTN increased ventilation and HCVR without any effect on metabolic rate^36^. Importantly, CO_2_ sensitive neurons do not contain MORs, which explains their resilience to opioids and attractiveness as a therapeutic target to excite downstream vulnerable respiratory neurons that are inhibited by opioids to treat OIRD^37,38^. A recent study by other investigators showed that optogenetic stimulation of *Phox2b+* RTN neurons activates breathing in conscious mice treated with fentanyl^51^. We have now shown that MC4R+RTN neurons project to respiratory premotor neurons in rVRG retrogradely labelled by the CTB administration to the phrenic nucleus and that both SET and photoexcitation of the MC4R+ RTN neurons induce excitatory neurotransmission to rVRG respiratory premotor neurons, which are opioid sensitive and MC4R- and project to the phrenic nucleus. All of the above suggest that SET treats OIRD by stimulating opioid-insensitive MC4R+ RTN neurons, which excite opioid-sensitive respiratory pre-motor neurons. Furthermore, SET does not bind to MOR, which explains the lack of effect on analgesia and amplify the potential for clinical use.

Our study has some limitations. First, although we have shown that SET attenuated OIRD induced by non-lethal doses of opioids we did not examine the effect of SET on opioid-mortality after administration of the lethal doses. Second, the dose of SET has been selected based on the previous metabolic literature^52–54^ and our recent study of SET in SBD^36^, but the dose response to SET has not been performed. Third, although our data suggest that SET may reverse OIRD acting in RTN, other mechanisms are possible. MC4R agonists may stimulate breathing in other brainstem centers controlling breathing, *e.g.* in other CO_2_ sensing areas such as dorsal and medullary raphe nuclei or in the *pre*-*Bötzinger* complex, Kölliker–Fuse nucleus, *etc.*, all of which show only low levels of MC4R expression^55,56^. Fourth, given that SET does not have affinity for MOR, we would expect that, unlike naloxone and nalmefene, SET would not induce opioid withdrawal, but it requires additional testing. Fifth, SET decreased apnea index in female mice, but more detailed exploration of sex differences in OIRD and MC4R signaling, as well as of the respiratory effects of MC4R agonists is needed.

In conclusion, our study provides the first evidence that the FDA approved for genetic obesity MC4R agonist setmelanotide is effective as a respiratory stimulant, which can reverse and treat OIRD in rodent models without affecting analgesia.

## Acknowledgements

This study was carried out with the support of the NIH (R01 HL174409, R01 HL128970, and R01 HL133100-S1), AHA (24CDA1270910). This study was financed, in part, by the São Paulo Research Foundation (FAPESP), Brazil. Process Number #2024/13971-9. A special thanks to Coordenação de Aperfeiçoamento de Pessoal de Nível Superior - Brazil (CAPES).

Figure 1, 3, 8, and graphical abstract were created with BioRender.com.

## Online Data Supplement

### Methods

The experiments were conducted on lean male C57BL/6J mice (#000664) aged 20 - 24 weeks, diet-induced obese (DIO) mice induced (#380050), and obese female C57BL/6J mice obtained from The Jackson Laboratory, lean Sprague Dawley rats and *Mc4r-Cre* mice (#008330). The animals were kept at the George Washington University Animal Facility and housed under a 12-hour light/dark cycle, at a controlled temperature of 29 °C, with unrestricted access to water and feed. The animals received a standard diet (3.0 kcal/g, 4.4% fat, 13% of calories from fat) or a high-fat diet (TD 03584, Teklad WI, 5.4 kcal/g, 35.2% fat, 58.4% of calories from fat) until DIO mice reached at least 40 g. All experimental procedures were approved by the George Washington University Animal Care and Use Committee (protocol A2023-014).

### Ventilation and respiration measurements – whole-body plethysmography (WBP)

Mice were placed in the WBP to measure breathing. A two-stage crossover study was conducted in which mice were randomly assigned to receive IP administration of the opioid (morphine at 10 mg/kg or fentanyl at 0.5 mg/kg) + vehicle with a one-week washout period, followed by IP administration of the opioid + SET (1 mg/kg) or vehicle. A whole-body plethysmography (WBP) system was adopted and configured as described in previous studies^1–3^ to measure respiratory and ventilatory parameters, such as respiratory rate, minute volume, number of apneas, and apnea index. The plethysmographic chamber was calibrated to generate reliable signals for both the mobilized tidal volume and airflow. A transducer (emka Technologies) is capable of filtering and interpreting the pressure oscillation signal inside the reference chamber as an electrical signal, and the continuous flow of air in the chamber is ensured by high-resistance elements and maintained by positive and negative pressure sources connected to flow regulators (Alicat Scientific). A slow leak was allowed to achieve an equilibrium between the reference chamber pressure and atmospheric pressure. Parameters such as the rectal temperature of the animals, ambient temperature, relative humidity, reference chamber temperature, and chamber gas constant were analyzed to calculate the current volume flow using the equation by Drorbaugh and Fenn^4^, using the injection of a known volume of air and the resulting deflection of the chamber. The signals captured were digitized at a rate of 1,000 Hz (sampling frequency per channel) and recorded using LabChart 7 Pro software (version 7.2, ADInstruments, Dunedin, NZ).

The animals were acclimated to the system for a period of 3 hours, one week before the start of experimental recordings for basal respiration assessment. Next, a dose of morphine or fentanyl was administered as described above, and the animals were placed inside the humidified plethysmographic chamber at 29°C for 15 minutes to record basal respiration. After this period, a dose of setmelanotide or vehicle was administered, and the total duration of respiration recordings was 2 hours.

### Tail flick test

The tail flick test was used to measure the latency of the animal’s tail reflex when exposed to a painful thermal stimulus by immersion in hot water and was conducted as described in previous studies^5,6^. Lean and DIO male mice were acclimated to the restraint tube for a period of 3 days before the experiment, and on the day of evaluation, they were immobilized for about 30 seconds in this acrylic tube, and the distal third of the tail was immersed in hot water at 50 ± 1°C. The time until tail withdrawal was analyzed to assess nociception. The maximum exposure time to hot water was set at 15 seconds to avoid possible tissue damage to the animal. Baseline latency was determined before the interventions and measured twice with a 5-minute interval between measurements. After baseline measurements, fentanyl (0.5 mg/kg) or morphine (10 mg/kg) IP was administered randomly and cross-over, and after 15 minutes, SET (1 mg/kg) or vehicle IP was administered. The latency time for tail withdrawal was recorded after 15, 30, 60, and 120 minutes after the application of SET or vehicle.

### Electrophysiological recordings

Animals were deeply anesthetized with isoflurane and perfused transcardially with a glycerol-based artificial cerebrospinal fluid (aCSF) that contained (in mM): 252 glycerol, 1.6 KCl, 1.2 NaH2PO4, 1.2 MgCl, 2.4 CaCl2, 26 NaHCO3, and 11 glucose. The brain was carefully removed, and 270-μm-thick coronal slices of the medulla that contained CTB-labeled pre-motor neurons as well as ChR2-EYFP-labeled MC4R neurons and fibers were obtained with a vibratome. The slices were transferred to a solution of the following composition (in mM) 110 N-methyl-d-glucamine (NMDG), 2.5 KCl, 1.2 NaH2PO4, 25 NaHCO3, 25 glucose, 0.5 CaCl2, and 10 mM MgSO4 equilibrated with 95% O2-5% CO2 (pH 7.4) at 34°C for 15 min.

The slices were then transferred from NMDG-based aCSF to a recording chamber, which allowed perfusion (5–10 ml/min) of aCSF at room temperature (25°C) containing (in mM) 125 NaCl, 3 KCl, 2 CaCl2, 26 NaHCO3, 5 glucose, and 5 4-(2-hydroxyethyl)-1-piperazineethanesulfonic acid (HEPES) equilibrated with 95% O2-5% CO2 (pH 7.4). Individual pre-motor neurons that project to the phrenic motor neurons in the ventral horn of the cervical spinal cord were identified in the rostral ventrolateral medulla (RVLM) by the presence of the retrograde fluorescent tracer CTB. To examine postsynaptic currents in these pre-motor neurons, patch pipettes (2.5–3.5 MΩ) were filled with a solution consisting of (in mM) 150 KCl, 2 MgCl2, 2 ethylene glycol-bis(β-aminoethyl ether)-N,N,N′,N′-tetraacetic acid (EGTA), 10 HEPES, and 2 Mg-ATP (pH 7.3).

Voltage clamp whole cell recordings were made at a holding potential of −80 mV with an Axopatch 200 B and pClamp 8 software (Axon Instruments, Union City, CA). At the end of the experiments AMPA and NMDA glutamate receptors were blocked by adding into aCSF 6-cyano-7-nitroquinoxaline-2,3-dione (CNQX, 25 μM) and D (−)-2amino-5-phosphopentanoic acid (AP5; 50 μM) respectively. Drugs used in this electrophysiological study were purchased from Sigma-Aldrich Chemical (St. Louis, MO). Selective photostimulation of ChR2-YFP-labeled MC4R fibers surrounding pre-motor neurons was performed by using a 473-nm blue laser light (CrystaLaser, Reno, NV). A series of consecutive single stimulations (3-ms duration, at 1 Hz) were applied to each pre-motor neuron. Laser light intensity was kept constant across all experiments at an output of 10 mW.

